# Maternal Immune Activation Alters Temporal Precision of Spike Generation of CA1 Pyramidal Neurons by Unbalancing GABAergic Inhibition in the Offspring

**DOI:** 10.1101/2024.02.19.581109

**Authors:** Ernesto Griego, Camila Cerna, Isabel Sollozo-Dupont, Marco Fuenzalida, Emilio J. Galván

**Affiliations:** Departamento de Farmacobiología, Cinvestav, Ciudad de México, México; Centro de Neurobiología y Fisiopatología Integrativa, Facultad de Ciencias, Universidad de Valparaíso, Valparaíso, Chile; Instituto Nacional de Cancerología, Ciudad de México, México; Centro de Investigaciones sobre el Envejecimiento, CIE-Cinvestav, Ciudad de México, México; Dominick P. Purpura Department of Neuroscience, Albert Einstein College of Medicine, New York, United States

**Keywords:** Maternal immune activation, Hippocampus, CA1 pyramidal cells, Synaptic integration, Behavioral impairments

## Abstract

Maternal immune activation (MIA) represents a risk factor for neuropsychiatric disorders associated with neurodevelopmental alterations. A growing body of evidence from rodents and non-human primates shows that MIA induced by viral or bacterial infections results in several neurobiological alterations in the offspring. These changes may play an important role in the pathophysiology of psychiatric disorders like schizophrenia and autism spectrum disorders, whose clinical features include impairments in cognitive processing and social performance. Such alterations are causally associated with the maternal inflammatory response to infection rather than with the infection itself. Previously, we reported that CA1 pyramidal neurons of mice exposed to MIA exhibit increased excitability accompanied by a reduction in dendritic complexity. However, potential alterations in cellular and synaptic rules that shape the neuronal computational properties of the offspring remain to be determined. In this study, using mice as subjects, we identified a series of cellular and synaptic alterations endured by CA1 pyramidal neurons of the dorsal hippocampus in a lipopolysaccharide-induced MIA model. Our data provide evidence that MIA reshapes the excitation-inhibition balance by decreasing the perisomatic GABAergic inhibition impinging on CA1 pyramidal neurons. These alterations yield a dysregulated amplification of the temporal and spatial synaptic integration. In addition, MIA-exposed offspring displayed social and anxiety-like abnormalities. Collectively, these findings contribute to the understanding of the cellular and synaptic alterations underlying the behavioral symptoms present in neurodevelopmental disorders associated with MIA.

**Highlights:**

- LPS injection during pregnancy (MIA) increases cytokine production and decreases litter size.
- MIA increases the temporal summation of EPSPs in hippocampal neurons.
- MIA alters spatial summation and increases the probability of action potential discharge.
- MIA alters the inhibitory/excitatory balance of CA1 pyramidal cells.
- MIA alters the expression of GAD-positive interneurons.
- MIA alters the performance of several behavioral tests in offspring.

## 1. Introduction

Prenatal environmental factors, including maternal immune activation (MIA), can trigger neurodevelopmental alterations that have been implicated in the etiology of neuropsychiatric disorders, including schizophrenia and autism spectrum disorders. Both epidemiological studies and experimental animal models have provided evidence for the molecular, cellular, synaptic, and behavioral alterations induced by viral or bacterial MIA (Brown & Meyer, 2018; Griego et al., 2022; Han et al., 2021; Kwon et al., 2021). Although MIA in humans does not always lead to neuropsychiatric illnesses in children, rodent models have allowed researchers to establish causal relationships that identify MIA as a risk factor that, in combination with other factors such as genetic composition and postnatal stress, can exponentially increase an offspring’s susceptibility to developing neurodevelopmental and psychiatric disorders (Choudhury & Lennox, 2021; Estes & McAllister, 2016; Meyer, 2019).

MIA has been shown to be particularly pernicious to the development of the medial temporal lobe, including the hippocampus and dentate gyrus (Couch et al., 2021; Hui et al., 2020; Makinodan et al., 2008), structures that play critical roles in learning and declarative memory formation and processing(Bird & Burgess, 2008; Squire, 1992; Tulving & Markowitsch, 1998). A growing body of evidence has demonstrated that MIA promotes changes in the excitability (Griego et al., 2022; Patrich et al., 2016), synaptic transmission (Cieślik et al., 2020; Nakagawa et al., 2020), and morphology (Griego et al., 2022; Pekala et al., 2021) of hippocampal neurons. In addition, alterations in hippocampus-dependent behaviors have also been documented (Ito et al., 2010; Savanthrapadian et al., 2013). Altogether, this set of alterations suggests that MIA leads to disturbances in the computational properties of the neural circuits of the hippocampus. Nonetheless, the harmful effects of bacterial-induced MIA in the cellular and synaptic factors that shape the dendritic integration of excitatory inputs remain virtually uncharted.

In this study, we explored the consequences of prenatal exposure to inflammation triggered by bacterial antigens in the temporal and spatial components of synaptic integration in CA1 pyramidal cells (CA1 PCs) of the offspring dorsal hippocampus. In addition to previously reported affectations in morphological and biophysical properties (Griego et al., 2022), we found that CA1 PCs from MIA-exposed offspring exhibit reduced cannabinoid-sensitive GABAergic inhibition, resulting in an altered temporal resolution of synaptic integration and facilitation to reach the threshold for action potential firing. These neurophysiological changes are accompanied by increased anxiety behaviors and deficits in social performance, both behavioral features of neuropsychiatric disorders associated with MIA.

Our findings shed light on cellular and synaptic alterations that occur early in brain development and that are of the utmost relevance for understanding the pathophysiology of MIA-associated mental illness in offspring.

## 2. Methods and materials

### 2.1 Animals

All the animal procedures were approved by the Ethics Committee for Animal Research of the Center for Research and Advanced Studies (protocol number 0063-13) and followed the guidelines of the Mexican Official Norm for the use and care of laboratory animals (NOM-062-ZOO-1999) that mandates minimization of suffering and reduction of the number of experimental animals. Our experimental procedures also complied with the National Institutes of Health (NIH) and Animal Research: Reporting of In Vivo Experiments (ARRIVE) guidelines for the use and care of experimental animals. Efforts were maximized to reduce to a minimum the number of animals used per experiment and, more importantly, any unnecessary animal pain and distress.

### 2.2. Maternal immune activation model

Female C57BL/6 mice were mated with breeding males at sexual maturity (7–10 weeks old). The estrous cycle was monitored by a vaginal smear, and the criteria for pregnancy confirmation was the presence of a vaginal plug or spermatozoa observed in the smear. Pregnant mice were injected with *Escherichia coli* O26:B6 LPS (500 µg/kg; Sigma, St. Louis, MO, USA) prepared in a sterile isotonic saline solution (NaCl 0.9%) on gestational day 15 (G15) to induce a systemic immune activation, as previously reported (Griego et al., 2022). G15 represents the second half of mouse gestation, which roughly corresponds to the second and third trimesters in humans when infections confer the highest risk of developing neurodevelopmental disorders (Lydholm et al., 2019; Watson et al., 2006). Next, dams were bred and housed at our institutional animal facility under controlled environmental conditions as follows: 12:12 h light:dark inverted cycle, room temperature and humidity (22 ± 1°C and 50 ± 5%, respectively), food and drinking water *ad libitum*, and periodic veterinary supervision. The day of birth was denominated postnatal day 0 (P0). The health of pups was assessed daily from P0 to weaning at P21, avoiding direct physical contact, after which they were housed in cages with three to four animals of the same sex each. All the experimental procedures were performed on male animals at P31 ± 3.

### 2.3. Monitoring body temperature by infrared thermometry in dams

Body surface temperature was measured using an infrared thermometer (FR-1DM1, Microlife, FL, USA), as previously reported (Griego et al., 2022; Kawakami et al., 2018). The body temperature was measured on the same location (abdominal surface) immediately before LPS or saline inoculation (t = 0) and monitored every 30 min up to 3 h.

### 2.4. Quantification of serum cytokine levels by ELISA in dams

Dams’ serum samples were obtained by centrifuging blood samples at 5,000 x g for 5 min (4°C) to remove blood cells. Serum levels of tumor necrosis factor (TNF)-α and interleukin (IL)-6 were quantified by sandwich enzyme-linked immunosorbent assay (ELISA) kits specific for each cytokine (Preprotech, Cranbury, NJ, USA), as previously reported (Griego et al., 2022).

### 2.5. Hippocampal slice preparation

Male offspring at P31 ± 3 were deeply anesthetized with sodium pentobarbital (50 mg/kg, intraperitoneal administration) and promptly decapitated. The brains were gently removed and placed into an icy sucrose-based solution with the following composition (in mM): 210 sucrose, 2.8 KCl, 2 MgSO_4_, 1.25 Na_2_HPO_4_, 25 NaHCO_3_, 1 MgCl_2_, 1 CaCl_2_, and 10 D-glucose, constantly bubbled with a 95% O_2_/5% CO_2_ carbogen mixture. The details for the obtention of sagittal brain slices have been described in previous publications (Griego et al., 2022; Márquez et al., 2023). Individual slices were transferred to a submerged chamber and continuously perfused with artificial cerebrospinal fluid (aCSF) at 3–4 mL/min with the help of a peristaltic pump (120S, Watson-Marlow, Wilmington, MA, USA). The composition of aCSF was as follows (in mM): 125 NaCl, 2.5 KCl, 1.25 Na_2_HPO_4_, 25 NaHCO_3_, 2 MgCl_2_, 2 CaCl_2_, and 10 D-glucose; pH = 7.3–7.4; osmolarity = 280–290 mOsm. The recording temperature was maintained at 30 ± 1°C with the help of a single-channel temperature controller (TC-324C, Warner Instruments, Hamden, CT, USA).

### 2.6. Electrophysiology

Dorsal hippocampal slices were visualized using infrared differential interference contrast optics adapted to an FN1 Eclipse microscope (Nikon Corporation, Minato, Tokyo, Japan). All the electrophysiology experiments were performed in CA1 PCs. CA1 PCs were identified on the basis of their shape, somatic position, and firing pattern. The patch pipettes were pulled from borosilicate glass using a micropipette puller (P97, Sutter Instruments, Novato, CA, USA). The pipette tips had a resistance of 4–6 MΩ when filled with an internal solution with the following composition (in mM): 135 K^+^-gluconate, 10 KCl, 5 NaCl, 1 EGTA, 10 HEPES, 2 Mg^2+^-ATP, 0.4 Na^+^-GTP, 10 phosphocreatine, and pH ≈ 7.20–7.28. Biocytin (0.4%) was routinely added to the intracellular solution for *post hoc* morphological reconstruction of the recorded neurons. Whole-cell patch-clamp recordings were performed using an Axopatch 200B amplifier (Molecular Devices, San José, CA, USA), digitized at a sampling rate of 10 kHz and filtered at 5 kHz with a high-resolution, low-noise digitizer Digidata 1322A (Axon Instruments, Palo Alto, CA, USA). Series resistance (_≈_10–40 MΩ) was monitored throughout all experiments, and cells that exhibited a significant change in series resistance (> 20%) were excluded from the analyses. Digital signals were acquired and analyzed offline with the help of pCLAMP 11.2 software (Molecular Devices).

### 2.7. Analysis of temporal summation

Concentric bipolar electrodes were placed in the *stratum radiatum* of CA1 (_≈_100–150 µm from the *stratum pyramidale*) to stimulate the Schaffer collaterals. Trains of 10 excitatory postsynaptic potentials (EPSPs) were recorded at frequencies of 20 or 50 Hz in current clamp (CC) mode. Temporal summation is expressed as the increase in synaptic depolarization occurring along a train and is calculated as

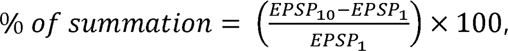

where EPSP_10_ is the peak amplitude of the tenth EPSP, and EPSP_1_ is the peak amplitude of the first EPSP in the train.

### 2.8. Analysis of spatial summation

Two concentric bipolar electrodes were placed in the *stratum radiatum* of CA1 located at _≈_100 and _≈_300 µm from the *stratum pyramidale* to stimulate two independent pathways of the Schaffer collaterals. Stimulation was delivered at time intervals (Δt) from 40 ms to 0 ms (steps of 5 ms), and vice versa, between the first and second stimulus. For each Δt, 10 trials were performed, and the probability of evoking an action potential (AP) due to the summation of EPSPs was calculated. Stimulation intensity was adjusted in such a way that, when the two Schaffer collateral pathways were stimulated simultaneously (Δt = 0 ms), the recorded neuron fired an AP in about 50% of the trials. Firing probability (P_Δt_) data was normalized and fitted to a Gaussian probability density function to obtain the standard deviation (SD), as previously reported (Pouille & Scanziani, 2001).

### 2.9. Analysis of spontaneous synaptic activity

Spontaneous excitatory and inhibitory postsynaptic currents (sEPSCs and sIPSCs, respectively) were recorded in voltage clamp (VC) mode, setting the holding potential (V_hold_) at −70 mV for sEPSC and 0 mV for sIPSC. sIPSCs were recorded in the presence of APV (50 µM) to block N-methyl-D-aspartate (NMDA) receptor-mediated transmission. At the end of the experiments, picrotoxin (PTX, 25 µM) was bath perfused to corroborate the GABAergic nature of the recorded events.

### 2.10. Analysis of GABAergic tonic current

GABAergic tonic current (I_tonic_) was assessed in VC mode as follows: Following a baseline recording of spontaneous activity (V_hold_ = 0 mV, in the presence of APV 50 µM), the selective GABA uptake inhibitor NO-711 (10 µM) was added to the bath to unveil the I_tonic_. Next, PTX (100 µM) was applied to the slice. The amplitude of I_tonic_ was measured as the difference in the holding current before and during the application of PTX.

### 2.11. Analysis of the cannabinoid receptor-1 and mu-receptor-sensitive GABAergic inhibition

Concentric bipolar electrodes were placed in the *stratum radiatum* of CA1 (_≈_50–100 µm from the *stratum pyramidale*) to stimulate GABAergic axons innervating CA1 PCs. Membrane potential was held at 0 mV, and APV (50 µM) was added to the bath perfusion to isolate the outward GABAergic currents. We took advantage of the selective expression of mu receptors (MOR) and cannabinoid receptor 1 (CB1R) in parvalbumin-expressing interneurons (PV-IN) and cholecystokinin-expressing interneurons (CCK-IN), respectively (Caccavano et al., 2024; Heifets et al., 2008; Monday et al., 2020) to analyze the effect of MIA on the selective inhibition from these two subpopulations of interneurons. Following a baseline recording, the selective MOR agonist DAMGO (2 µM) or the selective CB1R agonist WIN 55, 212 (2 µM) was added to the bath perfusion. PV-IN- and CCK-IN-dependent inhibition was calculated as the percentage decrease in IPSC amplitude in the presence of DAMGO or WIN relative to the baseline. At the end of the experiments, picrotoxin (PTX, 25-50 µM) was bath perfused to corroborate the GABAergic nature of the recorded events.

### 2.12. Immunofluorescence and confocal image acquisition

Male mice born to dams injected with LPS or saline were anesthetized with pentobarbital and intracardially perfused with phosphate-buffered saline (PBS 1X) followed by 4% paraformaldehyde (PFA) in a 0.1 M phosphate buffer (PB; pH 7.4). Brains were post-fixed in 4% PFA for 24 h at 4°C. Next, the brains were transferred to PBS 1X for at least 24 h. Coronal brain sections (50 μm) were obtained using a vibrating tissue slicer (Leica VT1000S; Nusschloc, Germany) and collected in culture plates filled with a cryoprotective solution (25% ethylene glycol, 25% glycerol, and 50% PB) and stored at −20°C until used. Immunolabeling was performed for glutamic acid decarboxylase 67 (GAD67). 4′,6-diamidino-2-phenylindole (DAPI) was included to visualize cell nuclei. Brain slices were washed with PBS and incubated with a blocking/permeation buffer (3% bovine serum albumin [BSA] and 0.3% Triton X-100 diluted in PBS) for 2–3 h at room temperature. Then, the slices were incubated overnight with a primary antibody (anti-GAD67, R&D Systems, mouse, MAB5406, AB_2278725) diluted at 1:500 in PBS with 2% BSA and 0.2% Triton X-100. Next, sections were washed and incubated for 2 h with a secondary antibody (anti-mouse Cy5, Innovative Research, 81-6516, AB_87836) diluted at 1:250 in PBS with 2% BSA and 0.2% Triton X-100. Sections were mounted using a Vectashield vibrance curing antifade mounting medium (Vector Laboratories, Burlingame, CA, USA; H-1800) and analyzed using a confocal microscope (LSM 800 with Airyscan; Carl Zeiss, Oberkochen, Germany). The frames (640 × 640 μm) for cell counting were selected randomly.

### 2.13. Analysis of offspring behavioral phenotype

All the behavioral tests were conducted at 31 ± 3 PN during the light cycle in the mice’s dimly lit testing room. Systematically, the animals were acclimatized to the behavior room at least 30 min before the test. A camera system filmed all the tests (acquisition at 30 FPS and 640 x 480 pixels, camera model C920, Logitech Co.). The video data were saved for offline analysis through a behavioral analysis program (ANY-maze v.7.09, Stoelting Co., Wood Dale, IL, USA).

### 2.14. Open-field test

Each mouse was individually placed in the center of a 40 x 40 cm square arena surrounded by high walls (30 cm) in a brightly lit room (300 lux), allowing free exploration of the open-field arena for 5 min (Fig. 6A), and a series of 10 x 10 cm zones (16 total blocks) were digitally defined using the analysis software. The outer zone consisted of 12 blocks, while the middle zone consisted of 4 blocks (20 x 20 cm). The analyses performed examined total distance traveled, average speed, maximum speed, time in the center, and time in the outer edge.

### 2.15. Elevated zero maze

The apparatus consisted of two open and two closed elevated arms (stress and protection arms, respectively) that formed a circle (height: 43.5 cm; diameter: 46 cm; lane width: 6 cm). Each mouse was individually placed in the open arms and allowed to freely explore the maze for 5 min (Fig. 7A). Anxiety levels were defined by the ratio of time spent in open arms to time spent in closed arms (Walf & Frye, 2007).

### 2.16. Sociability and social novelty preference test

A modified protocol of the three-chamber test described before was used (Crawley et al., 2007). The apparatus consisted of an open field with two containers. The first stage was habituation, in which the animal could move freely around the field for 5 min. The second stage was the sociability test, which consisted of placing a second animal (stranger I: S1) in one of the containers, and the test mouse explored for 10 min. The third stage was the social novelty preference, which consisted of placing a third animal (stranger II: S2) in the empty container while S1 remained in the same position (Fig. 8A). The test animal was allowed to explore for 10 min, and the time between each stage was set to 5 min. Social interaction occurred when the mouse nose-spoke, sniffed, or touched the container with the stranger. The time spent interacting in the first and second stages was quantified.

### 2.17. Statistical analyses

Results are expressed as mean ± SEM. The violin plots show the median, mean, interquartile range, and individual data. All statistical comparisons were performed between the offspring of saline-treated and LPS-injected dams. Animals were assigned randomly to experimental groups by simple random sampling, and the assignment was blinded to the experimenters to avoid any possible source of bias. For the electrophysiological data, the normality distribution of data was validated with the Kolmogorov–Smirnov test (*P* = 0.05). The comparability among experimental conditions was assessed by a two-tailed unpaired Student’s t-test or two-way ANOVA, as appropriate. When F achieved minimal statistical significance, the Holm– Šidák *post hoc* test was used for multiple comparisons. For all experiments, data were considered significant if *p* < 0.05. For behavioral data, a factor analysis was performed as previously described (Doremus et al., 2006). All test variables were checked for violations of normality. Skewness and kurtosis statistics were assessed, with a statistic of two times or less the standard error being considered acceptable for analysis. When a variable violated this criterion, the measure was subjected to transformation (e.g., square root, log (n+1), arc sine), using the transform to produce the most normal distribution for that variable. A principal components analysis was then conducted, with an orthogonal (varimax) rotation used on the factor matrix. Only components with an eigenvalue of one or greater were retained for final rotation. Additionally, the Kaiser–Meyer–Olkin (KMO) measure of sampling adequacy and Bartlett’s test of sphericity were run to ensure that these data were adequate for use in the analyses. Data analysis was performed with the SPSS software (SPSS Inc., Version 26, Chicago, IL, USA) and Clampfit 11.2 (Molecular Devices).

## 3. Results

### Intraperitoneal inoculation of LPS leads to sickness behavior and triggers the systemic release of pro-inflammatory cytokines in the dam

Pregnant mice (C57BL/6, 12 ± 2 weeks old) were inoculated with a single inoculation of LPS (500 µg/kg) on GD15 to induce MIA (Fig. 1A). In response to LPS, the dams exhibited signs of “sickness behavior” beginning at 0.5 h post inoculation and lasting up to 3 h. The sickness behavior included hypokinesia, piloerection, and a slight increase in body temperature, as previously reported (Biesmans et al., 2013; Griego et al., 2022). In addition, LPS-inoculated dams exhibited a slight increase in body temperature; however, this change was not statistically different compared to the control dams (Fig. 1B). In addition, dams that did not show signs of sickness behavior after LPS inoculation were excluded from the study. To corroborate the inflammatory response induced by LPS, levels of pro-inflammatory cytokines (TNF-α and IL-6) were determined in serum 1 h after inoculation with saline solution or LPS. As expected, LPS inoculation increased the serum levels of TNF-α (Student’s t-test; t_(8)_ = 16.320; *p* ≤ 0.001; n = 5 animals; Fig. 1C) and IL-6 (Student’s t-test; t_(8)_ = 9.119; *p* ≤ 0.001; n = 5 animals; Fig. 1D). The sickness behavior and elevated systemic levels of TNF-α and IL-6 confirm that systemic injection of *Escherichia coli* LPS is a potent stimulus to induce MIA in pregnant mice, as we previously reported using the same experimental protocol (Griego et al., 2022).

**Fig. 1.**
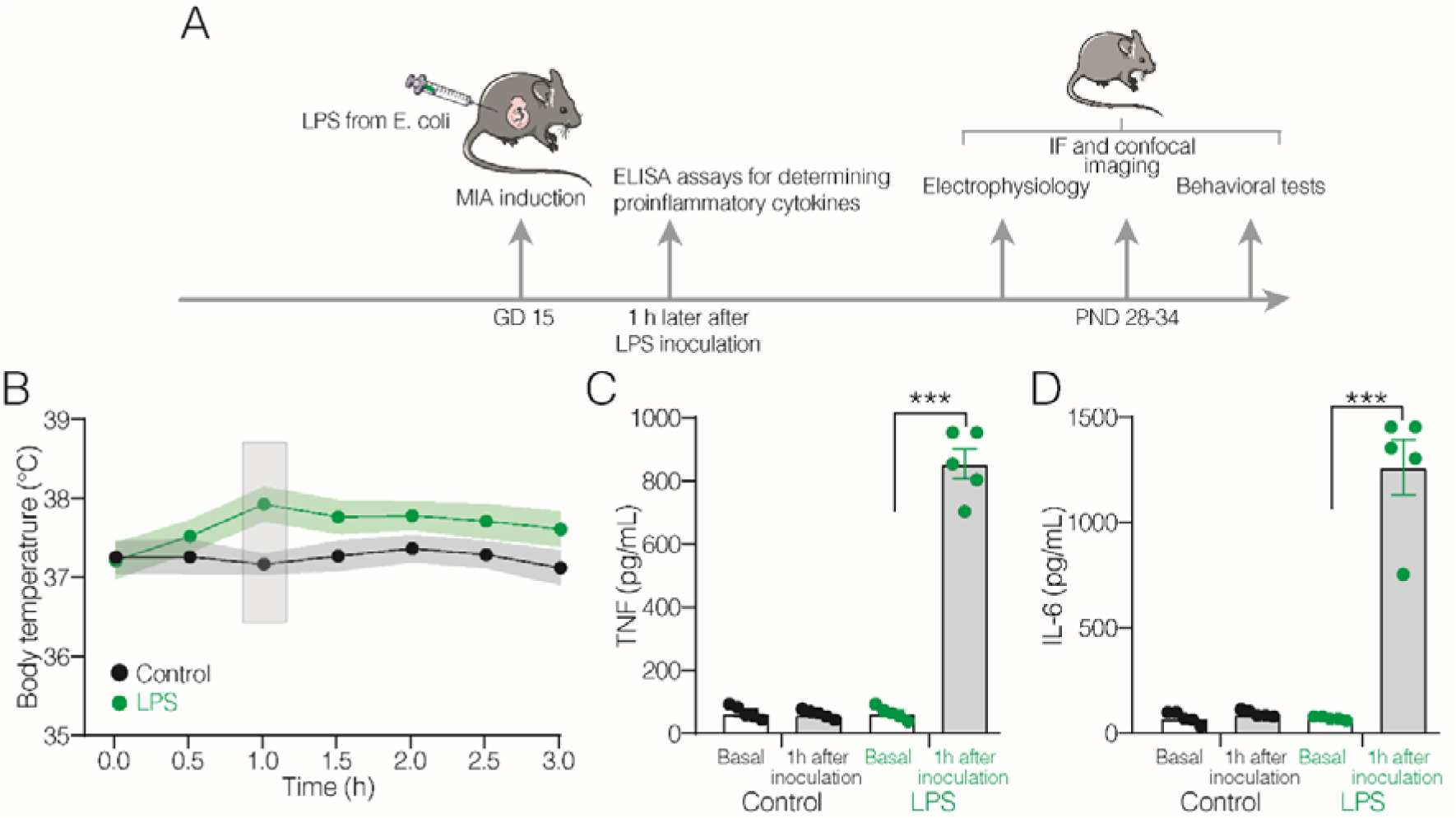
Intraperitoneal injection of LPS from Escherichia coli promotes MIA. **A)** Schematic representation of the experimental timeline. Pregnant C57BL/6 mice were inoculated with LPS on gestational day 15 (GD 15). ELISA assays were performed in dams to measure serum levels of pro-inflammatory cytokines. Patch-clamp recordings, immunofluorescence and confocal imaging evaluations, and behavioral experiments were performed in the offspring at postnatal day 28–35 (PND 28–35). **B)** Body temperature was measured before LPS or saline inoculation (time = 0) and every 30 min up to 3 h after inoculation. **C–D)** Bar graphs showing the levels of pro-inflammatory cytokines, TNF-α and IL-6, in maternal serum before and 1 h after LPS inoculation. Black symbols = control mice; green symbols = LPS-inoculated mice. \*\*\**P* < 0.001.

### LPS-induced MIA increases the temporal summation of dorsal CA1 PCs of the offspring

Previous studies have documented that MIA alters critical aspects of neurodevelopment, resulting in changes in neuronal morphology and function (Couch et al., 2021; Hanson et al., 2023; Savanthrapadian et al., 2013). In a previous study, we reported that LPS-induced MIA leads to increased intrinsic excitability and reduces the morphological complexity of dendrites of dorsal CA1 PCs of the offspring. We also reported that MIA causes an imbalance in the spontaneous GABAergic and glutamatergic activity impinging upon CA1 PCs (Griego et al., 2022). Since intrinsic excitability, geometric properties of dendrites, and GABAergic inhibition shape the computational capabilities of neurons (Magee, 2000; Pouille & Scanziani, 2001; Spruston, 2008), we tested the hypothesis that MIA alters the synaptic integration of the offsprings’ hippocampus. To test this idea, we examined the temporal and spatial components of synaptic integration of CA1 PCs in acute hippocampal slices. Fig. 2A1 shows a representative biocytin-filled CA1 PC included in the study. Temporal summation of synaptic activity was evaluated directly via extracellular stimulation of Schaffer collaterals (Fig. 2A2), and trains of excitatory postsynaptic potentials (EPSPs) were evoked at two physiologically relevant stimulation frequencies (20 and 50 Hz) (Combe et al., 2018; Marcelin et al., 2012; Masurkar et al., 2020) in control and MIA-exposed hippocampal slices (Fig. 2A2). The amplitude of the EPSPs, measured along the stimulation train, revealed a consecutive increase in the EPSP amplitude in both experimental conditions and the stimulation frequencies tested. At 20 Hz stimulation, no statistical difference was found in the percentage of summation of MIA-exposed CA1 PCs compared to the control cells (summation in control: 69.37 ± 1.86%; in MIA: 72.47 ± 1.63%; Student’s t-test; t_28_ = 1.253; *p* = 0.220; n = 15 cells / 10 animals / 5 litters for each condition; Supp Fig. S1). By contrast, the summation elicited at 50 Hz stimulation revealed a significant response in the MIA-exposed CA1 PCs compared to the control cells (summation in control: 201.91 ± 5.60%; in MIA: 312.60 ± 6.74%; Student’s t-test; t_28_ = 12.620; *p* < 0.001; n = 15 cells / 10 animals / 5 litters for each condition; Fig. 2B-D). These results indicate that MIA-exposed cells exhibit a larger capability to sum excitatory inputs originating from the same presynaptic origin.

**Fig. 2.**
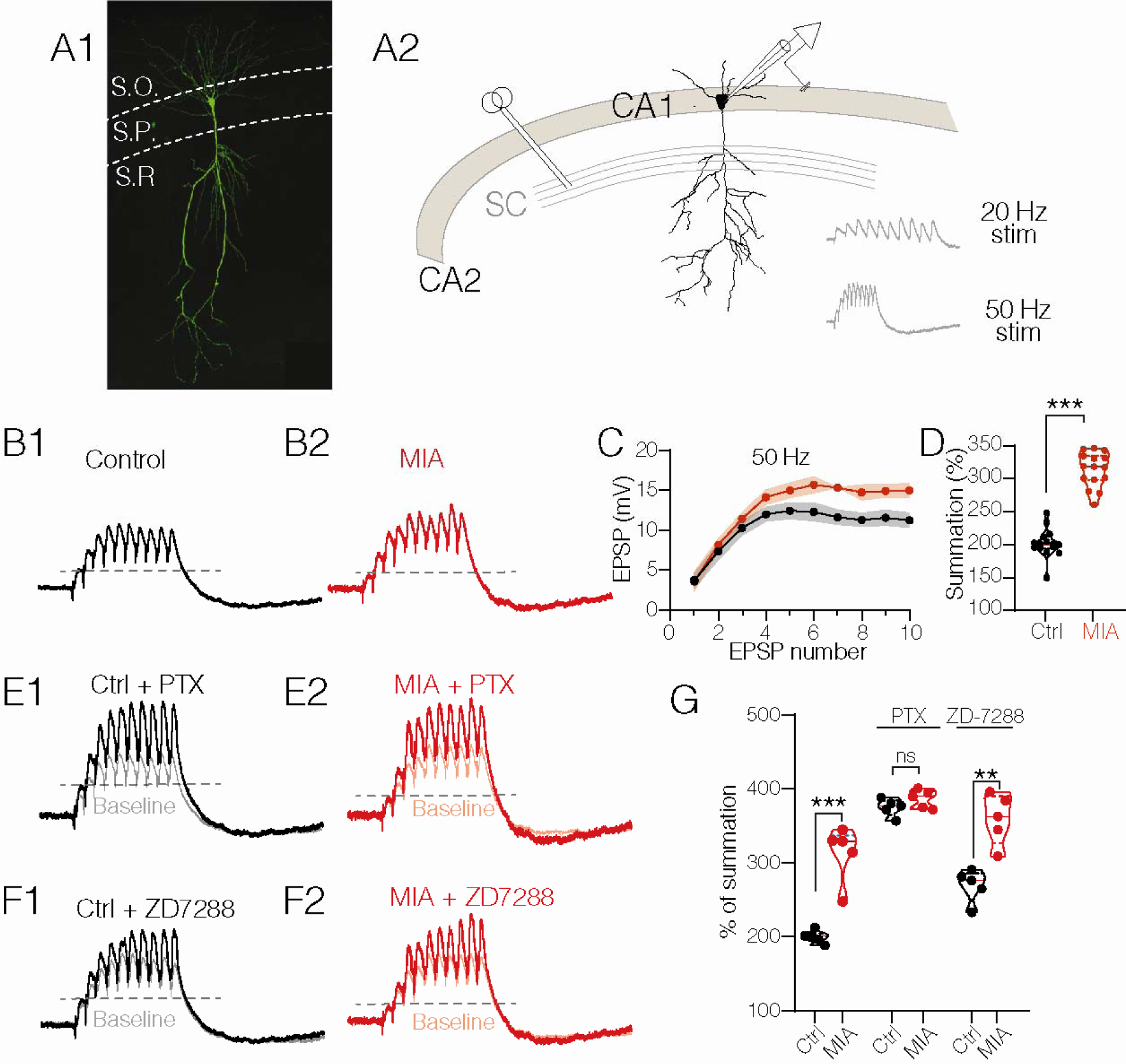
MIA alters the temporal summation of CA1 PC of the offspring. **A1)** Confocal image of a representative biocytin-filled CA1 PC included in the study. **A2)** Scheme of the electrode positioning. Temporal summation was evoked with stimulation trains delivered at 20 and 50 Hz. **B1, B2)** Representative traces of the EPSP trains at 50 Hz in both experimental conditions. **C)** Scatter plot showing EPSP amplitude along the 50 Hz train evoked by Schaffer collaterals stimulation. **D)** Violin plot summarizing the effect of MIA on the temporal summation of EPSPs evoked at 50 Hz. **E1, E2)** Representative traces elicited at 50 Hz in both experimental conditions after a bath perfusion of picrotoxin (PTX, 50 µM). **F1, F2)** Representative traces elicited at 50 Hz after a bath perfusion of ZD7288 (10 µM). **G)** Violin plots summarizing the change in temporal summation under the different pharmacological conditions. Although ZD7288 increased the percentage of summation in both experimental conditions, it failed to equalize the summation in both groups, whereas PTX increased the percentage of summation and equalized it. \*\**P* < 0.01; \*\*\**P* < 0.001.

To test the hypothesis that the increased temporal summation of MIA-exposed animals may result from an impairment in their GABAergic transmission, we added PTX (50 µM) to the bath to block the GABA_A_-receptor-mediated synaptic inhibition. Fig. 2E shows superimposed traces of EPSPs evoked at 50 Hz in the control and following PTX perfusion. As expected, PTX increased the temporal summation, equalizing the percentage of summation in both control and MIA-exposed mice (summation in control: 374.13 ± 5.27 %; in MIA: 385.74 ± 5.55 %; Student’s t-test; t_8_ = 1.504; *p* = 0.171; n = 5 cells / 3 animals / 2 litters for each condition; Fig. 2G). In area CA1, temporal summation depends on the dendritic conductances mediated by hyperpolarization-activated cyclic nucleotide-gated channels (HCN) or *h*-conductance, and its blockade with ZD-7288 suppresses temporal summation (Magee, 1999). To test the involvement of the *h*-conductance in the increased temporal summation of the MIA-exposed cells, ZD-7288 (20 µM) was bath perfused in another group of CA1 PCs. ZD-7288 effectively increased the percentage of summation evoked at 50 Hz in both experimental conditions (summation in control: 268.31 ± 9.96%; in MIA: 358.4 ± 15.38%; Student’s t-test; t8 = 4.916; *p* = 0.001; n = 5 cells / 3 animals / 2 litters for each condition; Fig. 2G). However, the statistical difference in the percentage of temporal summation between the experimental groups persisted (right-side violins, Fig. 2G), suggesting that the *h*-conductance does not participate in the increased temporal summation of MIA-exposed CA1 PCs.

### LPS-induced MIA increases the spatial summation of dorsal CA1 PCs of the offspring

Next, we investigated possible changes in the spatial summation of excitatory inputs onto CA1 PCs. For this purpose, two extracellular stimulation electrodes (S1 and S2; see the schematic representation, left panel in Fig. 3A) were placed in the *stratum radiatum,* 150 to 400 µm away from the *stratum pyramidale,* with an interelectrode distance of _≈_125 ± 25 µm. The intensity of the electrical stimulation was then adjusted to a level at which, with simultaneous delivery of the two stimuli, the neuron evoked an action potential in approximately 50% of the trials. The right panel in Fig. 3A shows EPSPs under the double stimuli protocol. The stimuli were delivered in time intervals of 5 ms, and the distribution of firing probabilities (P_Δt_) against each time interval (Δt) was normalized and plotted. A Gaussian function was fitted to the resulting histograms to calculate the standard deviation of distribution (SD) as a measure of integration window narrowness. The resulting histogram of the P_Δt_ in the control condition is depicted in Fig. 3B (n = 15 cells / 10 animals / 5 litters). Because GABAergic inhibition exerts a powerful control over spatial summation and the integration of excitatory inputs (Pouille & Scanziani, 2001), we analyzed the effect of suppressing the GABAergic inhibition on the P_Δt_. Fig. 3C shows the histogram of the firing probabilities of the control CA1 PCs in the presence of picrotoxin (PTX, 50 µM). As expected, we found a significant increase in the SD of the integration window (SD in control: 7.28 ± 0.35 ms; in control + PTX: 21.34 ± 0.70 ms; Student’s t-test; t_18_ = 18.64; *p* < 0.001; Fig. 3G). The inset graph in Fig. 3C compares the narrowness of the integration window of CA1 PCs in the control and the presence of PTX.

**Fig. 3.**
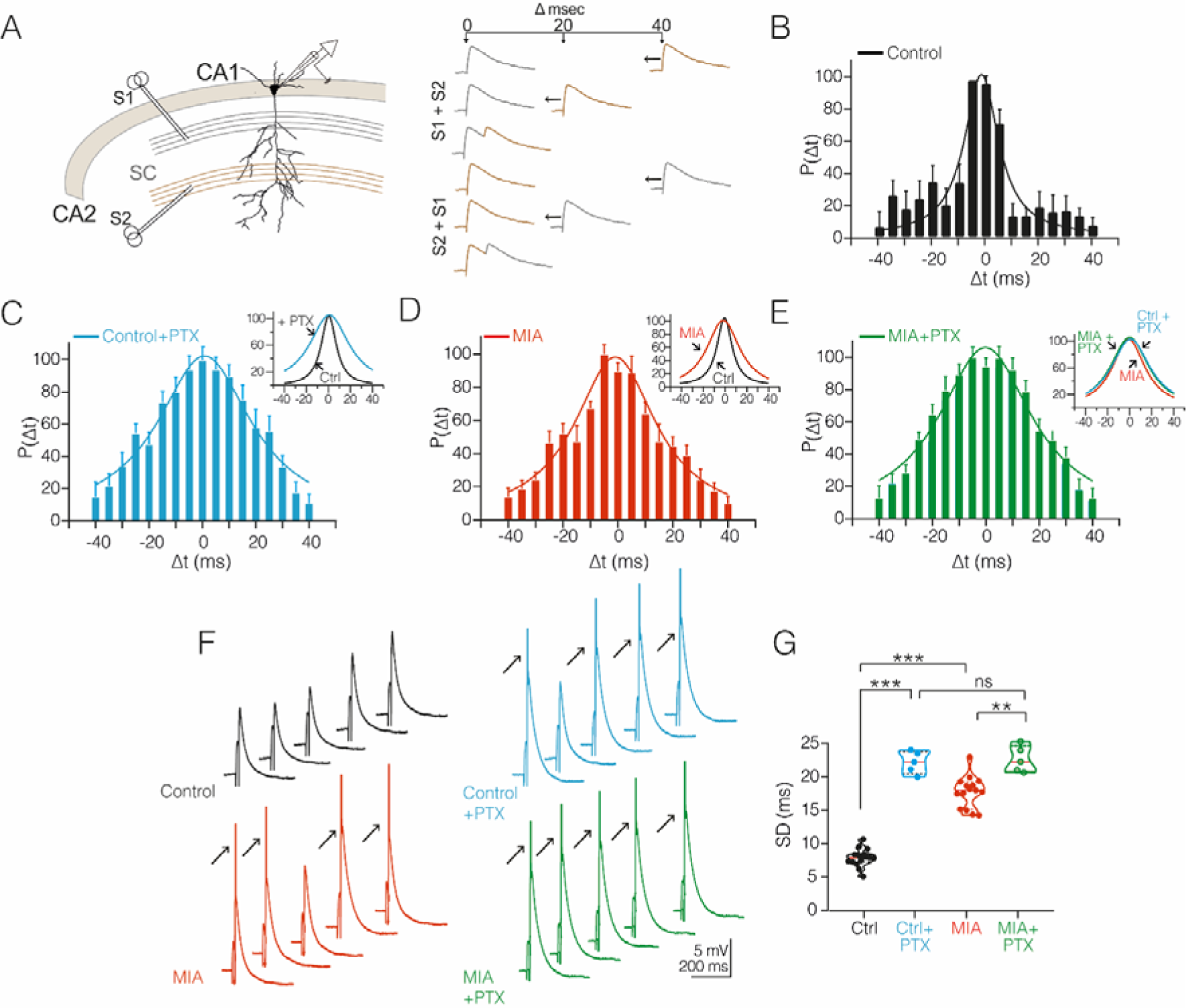
MIA alters the spatial summation of CA1 PC of the offspring. **A)** Electrode positioning for the electrophysiologic recordings in acute hippocampal slices. Spatial summation was achieved by stimulating two independent inputs from the Schaffer collaterals at different interstimulus intervals (Δt, from 40 ms to 0 and vice versa in increments of 5 ms). **B–E)** PΔt data histograms at the different Δt evaluated in control (B), control +PTX condition (C), MIA (D), and MIA +PTX (E). Standard deviation was calculated by fitting a Gaussian probability density function to each histogram. **F)** Representative voltage traces showing spatial summation of EPSPs at Δt = 20 ms in all the experimental conditions. Arrowheads indicate the evoked spikes (truncated trace) when the EPSPs sum reached the firing threshold. **G)** Violin plots summarizing the effect of MIA and the blockade of GABAergic transmission on the integration window standard deviation. **P<0.01; ***P<0.001.

Next, we analyzed the spatial summation in MIA-exposed mice. Fig. 3D shows the Gaussian function fitted to histograms of P_Δt_, revealing a significant increase in the SD of the integration window of MIA-exposed CA1 PCs compared to the control cells (SD in MIA: 17.71 ± 0.61 ms; Student’s t-test; t_28_ = 14.160; *p* < 0.001; n = 15 cells / 10 animals / 5 litters; Fig. 3G). The inset graph in Fig. 3D contrasts the narrowness difference of the P_Δt_ of the control cells and MIA-exposed cells. Interestingly, as shown in Fig. 3E, the blockade of GABAergic inhibition with PTX slightly increased the P_Δt_ of MIA-exposed CA1 PCs (SD in MIA +PTX: 21.94 ± 0.84 ms; Student’s t-test; t_18_ = 3.855; *p* = 0.002; Fig. 3G). Moreover, the blockade of inhibition resulted in the unwinding of the differences in P_Δt_ between both the control and MIA-exposed neurons (Control + PTX vs MIA + PTX; Student’s t-test; t_10_ = 0.597; *p* = 0.563; Fig. 3G). Fig. 3F shows representative voltage traces of EPSPs evoked in the different experimental conditions at Δt = 20 ms and the resulting action potential (arrows) when synaptic stimulation reached the spike threshold. Collectively, these results strongly suggest that LPS-induced MIA promotes an imbalance in GABAergic transmission of area CA1 that alters the temporal fidelity of spikes and the synaptic integration capabilities of CA1 PCs.

### LPS-induced MIA imbalances the excitatory-inhibitory ratio in CA1 PCs of the offspring

The integrative skills of CA1 PCs, including spatial and temporal summation, are controlled by several factors, such as intrinsic excitability and the geometric properties of their dendritic trees (Magee, 2000; Spruston, 2008). In line with this tenet, local interneurons are known to regulate these capabilities (Pouille & Scanziani, 2001). Therefore, the next experiments aimed to confirm the possible imbalance in GABAergic inhibition by evaluating molecular and functional markers of GABAergic transmission in the MIA-exposed offspring. For this purpose, we quantified the expression of GAD-67, the limiting enzyme for GABA synthesis, in area CA1 of the dorsal hippocampus. Fig. 4A shows representative confocal microscopy images showing the reduction in the number of GAD-67 positive cells in MIA-exposed animals compared to the control animals (GAD-67-expressing cells per field in control: 60.56 ± 4.45 cells; in MIA: 45.44 ± 5.48 cells; Student’s t-test; t_16_ = 2.139; *p* = 0.048; n = 3 animals / 3 slices per animal for each condition; Fig. 4B). These results strongly suggest either a loss of GABAergic interneurons or a reduction in the expression of GAD-67 by local CA1 interneurons. In either of these possible scenarios, our data strongly indicate a reduction in the GABAergic strength of the hippocampal network. To further explore this possibility, we recorded spontaneous synaptic activity from the CA1 PCs of the offspring. Fig. 4C shows representative traces of spontaneous excitatory and inhibitory postsynaptic currents (sEPSCs and sIPSCs, respectively) in both the control and MIA conditions. sEPSCs were recorded holding the membrane potential at −70 mV, whereas sIPSCs were recorded at 0 mV in the presence of AP-V (25 µM) to suppress the NMDA-receptor-dependent outward currents. Statistical analysis of synaptic activity revealed that MIA slightly increased the amplitude and frequency of sEPSCs (sEPSC amplitude in control: 13.51 ± 0.82 pA; in MIA: 16.45 ± 1.16 pA; Student’s t-test; t_28_ = 2.054; *p* = 0.049; sEPSC frequency in control: 9.04 ± 0.65 Hz; in MIA: 11.18 ± 0.79 Hz; Student’s t-test; t_28_ = 2.088; *p* = 0.046; n = 15 cells / 10 animals / 5 litters for each condition; Fig. 4D1). On the other hand, the amplitude of the sIPSCs of MIA-exposed mice did not exhibit significant changes (sIPSC amplitude in control: 30.13 ± 1.91 pA; in MIA: 26.89 ± 2.31 pA; Student’s t-test; t_28_ = 1.081; *p* = 0.288; n = 15 cells / 10 animals / 5 litters for each condition; Fig. 4D2). However, the frequency of sIPSCs was dramatically decreased in the MIA-exposed mice, suggesting an alteration of the GABA release process at the presynaptic level (sIPSC frequency in control: 5.04 ± 0.24 Hz; in MIA: 3.31 ± 0.25 Hz; Student’s t-test; t_28_ = 4.910; *p* < 0.001; Fig. 4D2). Next, we explored potential changes in the excitatory-inhibitory (E-I) balance of CA1 PCs of the offspring by evoking EPSCs and IPSCs via electrical extracellular stimulation. EPSCs were recorded holding the membrane potential at −70 mV. It should be noted that, at this holding potential, NMDA receptors are blocked by Mg^2+^ ions. Consequently, the recorded current is predominantly mediated by AMPA receptors. IPSCs were recorded at 0 mV in the presence of AP-V (25 µM) and CNQX (50 µM) to suppress NMDA- and AMPA-receptor-dependent currents. At the end of the experiment, the GABA_A_ receptor antagonist picrotoxin (PTX, 50 µM) was bath perfused to corroborate the GABAergic nature of the recorded outward currents. Fig. 4E shows representative traces recorded from the control and MIA-exposed CA1 PCs. No differences were found in the EPSCs’ amplitude between the control and MIA-exposed mice (bottom traces, Fig. 4E). However, the amplitude of the IPSCs of the MIA-exposed cells exhibited a significant decrease compared to the control cells (upper traces, Fig. 4E). Accordingly, our data reveal that MIA alters the relationship between EPSCs and IPSCs’ amplitude (slope of linear fit in control: 1.12; in MIA: 0.59; n = 15 cells / 10 animals / 5 litters for each condition; Fig. 4F). As a consequence, the E-I balance exhibited a shift toward excitation (E-I ratio in control: 0.89 ± 0.03; in MIA: 1.11 ± 0.04; Student’s t-test; t_28_ = 3.861; *p* = < 0.001; Fig. 4G).

**Fig. 4.**
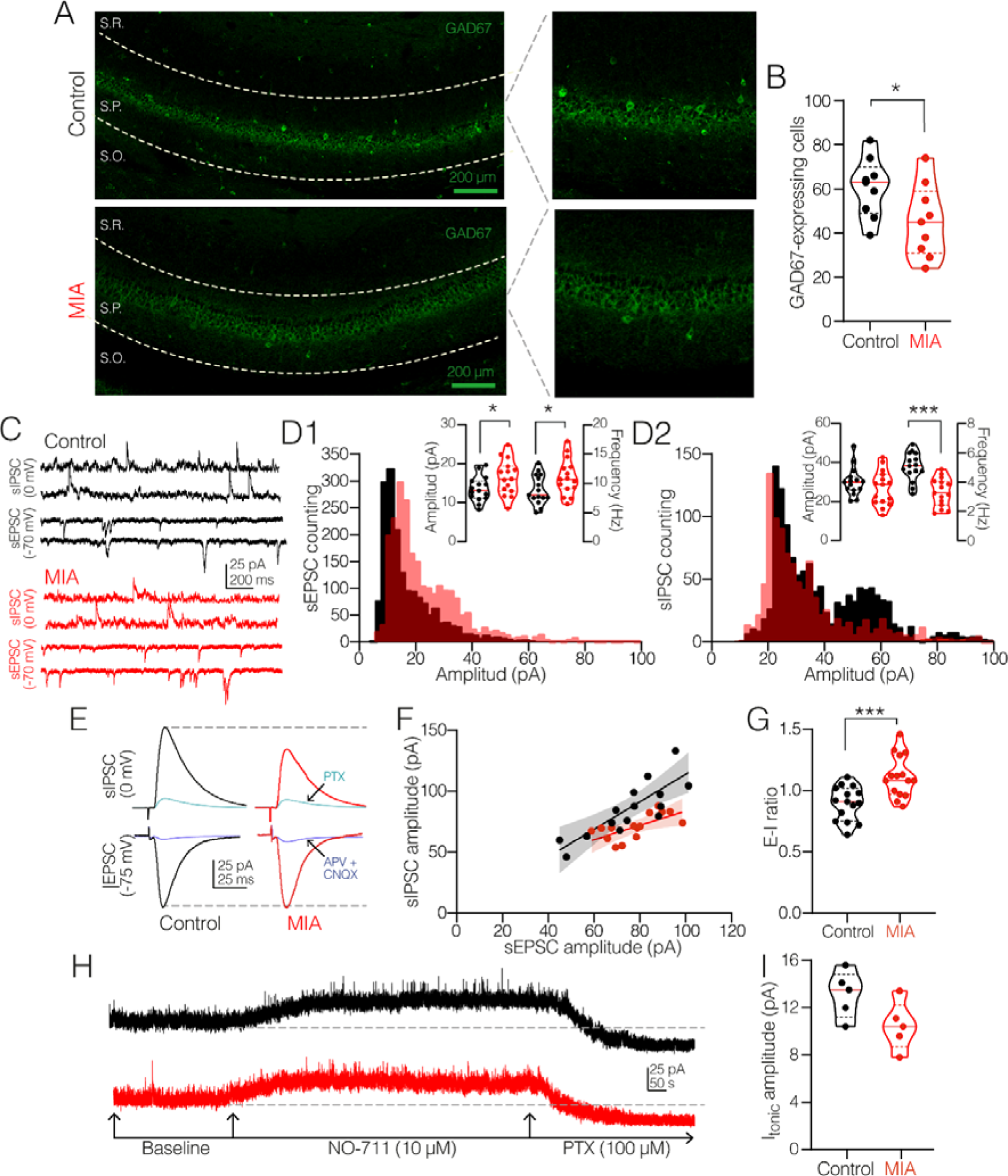
MIA alters the E-I balance in CA1 PC of the offspring. **A)** Representative confocal images showing a reduction in the number of interneurons immunoreactive to GAD-67 in the CA1 hippocampal area of the MIA-exposed offspring. **B)** Violin plots summarizing the effects of MIA on the density of CA1 GABAergic interneurons. Each dot represents the average of three counting frames per slice. n = 9 slices / 3 animals / 3 litters for each condition. **C)** Representative traces of spontaneous synaptic activity recorded in the CA1 PC. sEPSCs and sIPSCs were recorded at −70 mV and 0 mV, respectively. **D1)** Histogram of the distribution of sEPSC amplitudes. Inset, violin plots summarize the effect of MIA on the amplitude and frequency of sEPSC in the CA1 PC of the offspring. **D2)** Histogram of the distribution of sIPSC amplitudes. Inset, violin plots summarizing the effect of MIA on the amplitude and frequency of sIPSC in the CA1 PC of the offspring. **E)** Representative traces of electrically evoked EPSCs and IPSCs in the CA1 PC. EPSCs and IPSCs were sensitive to glutamatergic and GABAergic blockers, respectively. **F)** Averaged EPSC amplitudes plotted against averaged IPSC amplitudes. Lines = linear regression fits. **G)** Violin plots showing that the excitation-inhibition (E-I) ratio was significantly increased in the CA1 PC from MIA-exposed offspring. n = 15 cells / 10 animals / 5 litters for each condition. **H)** Representative traces of tonic GABAergic currents (I_tonic_) recorded in the CA1 PC from both the control and MIA-exposed offspring and (bottom) the pharmacological tools used for its measurement. **I)** Violin plots showing that I_tonic_ was not significantly affected by MIA in the CA1 PC of the offspring. Black symbols/traces = control cells; red symbols/traces = MIA cells. n = 5 cells / 4 animals / 2 litters for each condition. \**P* < 0.05; \*\*\**P* < 0.001.

Previous studies have documented two distinct forms of GABAergic inhibition impinging on CA1 PCs: a low-amplitude, slow-kinetic tonic current and a fast, phasic synaptic current (Domínguez et al., 2016; Gasulla & Calvo, 2015). What is more, tonic GABA_A_-receptor-mediated currents provide an important restraint on the excitability of CA1 PCs (Bonin et al., 2007; Kammel et al., 2018). To explore whether tonic inhibition was also mediating the MIA-related inhibition imbalance, we recorded CA1 PCs under the same conditions as for spontaneous inhibitory activity but added the GABA transporter inhibitor NO-711 (10 µM) in the bath. This pharmacologic strategy allowed us to unmask GABAergic tonic currents (*I*_tonic_) by increasing the extrasynaptic GABA concentration. After an NO-711 perfusion, PTX (100 µM) was added to the bath to block both phasic and tonic GABAergic currents. Fig. 4H shows representative traces of the GABAergic *I*_tonic_ recorded using this pharmacological approach. Nevertheless, although GABAergic *I*_tonic_ in MIA-exposed CA1 PCs displayed a slightly decreased amplitude, this change did not reach statistical significance compared to the control cells (*I*_tonic_ amplitude in control: 13.12 ± 0.89 pA; in MIA: 10.46 ± 0.91 pA; Student’s t-test; t_8_ = 2.078; *p* = 0.071; n = 5 cells / 3 animals / 2 litters for each condition; Fig. 4I). Taken together, these results strongly suggest that LPS-induced MIA dramatically impacts GABAergic inhibition, promoting a tendency toward hyperexcitability that may explain the higher synaptic summation displayed by the CA1 PCs of the offspring.

### MIA alters the perisomatic inhibition onto CA1 pyramidal cells

Synaptic integration of excitatory inputs is critically regulated by perisomatic inhibition. This inhibitory restraint is predominantly controlled by interneurons that express the calcium-binding protein parvalbumin and the neuropeptide cholecystokinin (PV-IN and CCK-IN, respectively) (Chevaleyre & Piskorowski, 2014; Freund & Katona, 2007). On the other hand, previous studies have documented dysregulation in the expression of GABAergic interneurons in response to MIA (Canetta et al., 2016; Vasistha et al., 2020). Therefore, in the next series of experiments, we used a pharmacological approach to investigate whether MIA alters the perisomatic inhibition mediated by PV-IN and CCK-IN. IPSCs were recorded in the CA1 PCs of the control and MIA-exposed mice by stimulating the axons of local interneurons in the presence of the glutamate blockers AP-V and CNQX (25 µM and 50 µM, respectively) and holding the membrane potential of the recorded PC at 0 mV. Fig. 5A1 shows a diagram illustrating the recording configuration and the sensitivity to PTX of the evoked synaptic response. After 10 min of baseline recording, the cannabinoid receptor type 1 (CB_1_R) agonist WIN55, 212-2 (1 µM), or the μ-opioid receptor (MOR) agonist, DAMGO (1 µM), was bath perfused. Since CB_1_R is mainly expressed in the CCK-IN and MOR is expressed in the PV-IN (Drake & Milner, 2002; Freund & Katona, 2007; Marsicano & Lutz, 1999), this pharmacological strategy was used to block GABA release in an interneuron-specific manner. The timeline and drugs employed in these experiments are summarized in Fig. 5A2. Figs. 5B1 and 5B2 show representative traces acquired in both experimental conditions, and the time-course graph shows the effect of DAMGO on the amplitude of the IPSCs, respectively. In the control cells, the bath perfusion of DAMGO reduced two-fold the amplitude of the IPSCs. In the MIA-exposed neurons, the DAMGO-mediated inhibition was slightly decreased; however, it did not reach statistical significance compared to the control cells (IPSC amplitude in control: 56.34 ± 3.17 % of the baseline; in MIA: 66.85 ± 3.67 % of the baseline; Student’s t-test; t_9_ = 1.840; *p* = 0.099; n = 10 cells / 5 animals / 3 litters for each condition; Fig. 5C). Likewise, the effect of WIN in the amplitude of the IPSCs is depicted in the representative traces of Fig. 5D1 and the time-course graph of Fig. 5D2, respectively. The bath perfusion of WIN decreased the IPSCs’ amplitude in the control cells; more importantly, this effect was significantly occluded in the MIA-exposed CA1 PCs (IPSC amplitude in control: 71.88 ± 1.57 % of the baseline; in MIA: 87.71 ± 2.14 % of the baseline; Student’s t-test; t_9_ = 4.893; *p* < 0.001; n = 10 cells / 5 animals / 3 litters for each condition; Fig. 5E). These results strongly suggest that GABAergic inhibition mediated by CB_1_R- and MOR-expressing perisomatic interneurons (CCK-IN and PV-IN, respectively) is altered in the hippocampus of animals exposed to MIA. Nonetheless, whether the lower effect of WIN involves alterations in the expression or function of CB_1_R requires additional investigation.

**Fig. 5.**
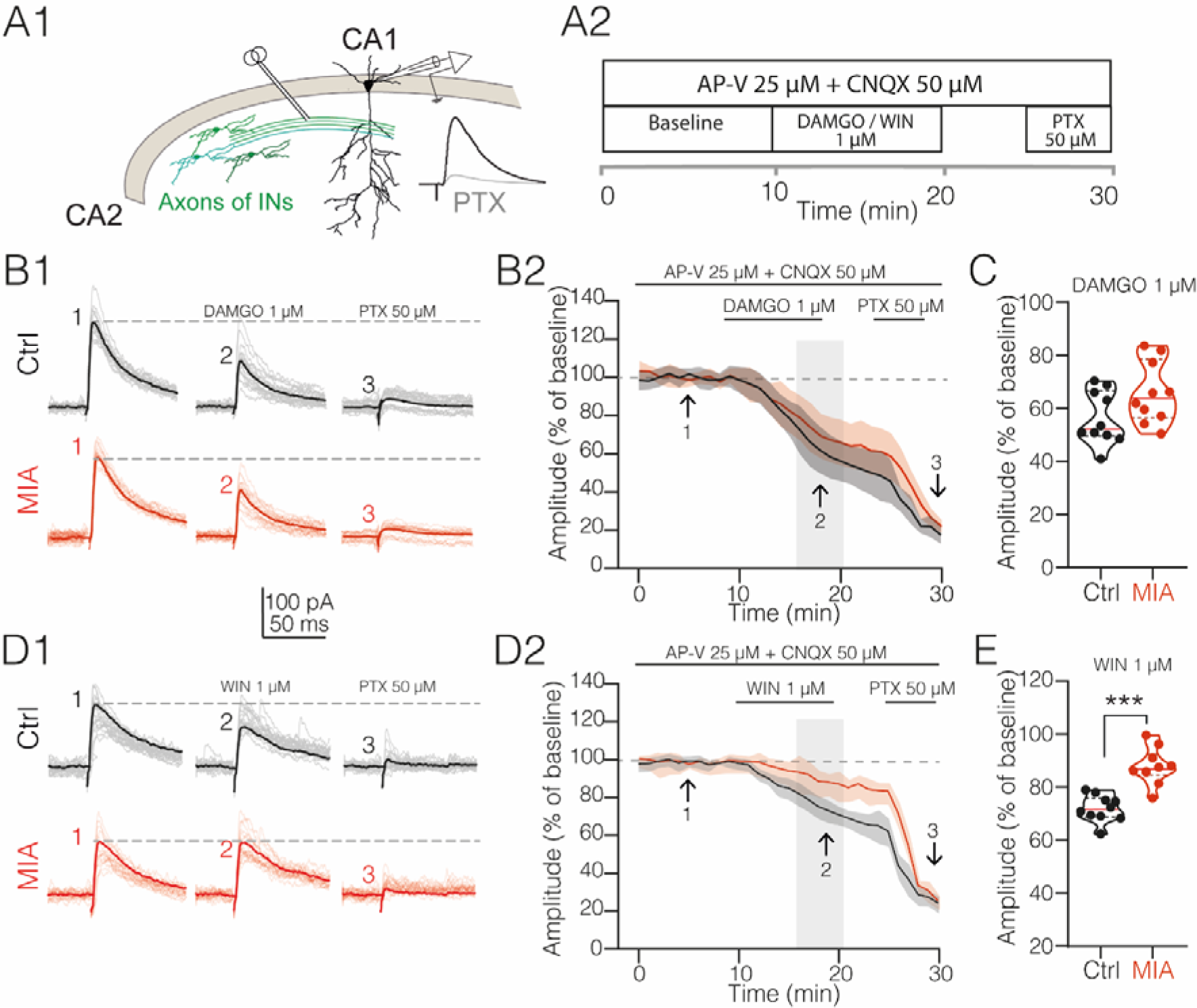
MIA affects the cannabinoid-sensitive inhibitory inputs in the CA1 PC of the offspring. **A1)** Schematic representation of the electrode positioning for electrophysiologic recordings. The GABAergic nature of the recorded IPSC was corroborated by a bath perfusion of PTX (50 µM) at the end of each experiment. **A2)** Table showing the experimental timeline and the pharmacological tools used for these experiments. **B1)** Representative IPSCs recorded in the CA1 PC from both the control and MIA-exposed offspring before (1) and after (2) DAMGO treatment (1 µM). **B2)** Summary time-course plot showing DAMGO-induced depression. The arrowhead indicates the time at which synaptic transmission was analyzed with respect to the baseline. **C)** Violin plots summarizing the reduction in the IPSC amplitude after DAMGO treatment. No difference was found between the control and MIA-exposed CA1 PC. **D1)** Representative IPSCs recorded in the CA1 PC from both the control and MIA-exposed offspring before (1) and after (2) WIN treatment (1 µM). **D2)** Summary time-course plot showing WIN-induced depression. The arrow indicates when plasticity was analyzed with respect to the baseline. **E)** Violin plots summarizing the depression in IPSC amplitude after WIN treatment. WIN-induced depression magnitude was significantly reduced in the CA1 PC from MIA-exposed mice. Black symbols/traces = control cells; red symbols/traces = MIA cells. n = 10 cells / 5 animals / 3 litters for each condition. \*\*\**P* < 0.001.

### MIA-exposed mice exhibit anxious-like behavior

For the next section, we determined possible changes in the locomotor activity, anxiety-like behavior, and social withdrawal in MIA-exposed mice since anxiety is often observed in individuals with schizophrenia and social withdrawal is categorized as a negative symptom of schizophrenia (Chamera et al., 2020). For this, we applied a principal component (PC) analysis stemming from the individual elements that compose these behaviors. The individual elements extracted and analyzed from each behavioral test are totaled in Table 1.

In the open-field test (OFT), the factor analysis yielded two principal components (PC1 and PC2, respectively) that explained 81% of the data set variance for the controls and 83% for the MIA-exposed mice (Table 1.1). In the control condition, the PC1 represented 43.5% of the total variance, whereas the MIA-exposed mice represented 44.9% (Figs. 6B and 6E, respectively). In both groups, PC1 was mainly carried by two measurements related to motor activity: distance traveled (DT) and average speed (AS) (Table 1.1). PC2 accounted for 38.1% of the variation in the two experimental groups (Figs. 6B and 6E, respectively). Time in the center (TC) and time in the outer edge (TOE) loaded highly on PC2 (Table 1.1); crucially, these variables (TC and TOE) are linked with anxiety-like behaviors. The comparison of the PCs strongly suggests that both groups had similar motor activity and anxiety levels.

In the elevated zero maze (EZM), the factor analysis yielded a result that accounted for 88.5% of the variance for the controls (Table 1.2 and Fig. 7B). This factor was constituted by most of the variables measured during the test, including time in the closed zone (TCZ), distance traveled (DT), average speed (AS), percentage of time in the opened zone (%OZ), and time in the opened zone (TOZ) (Fig. 7C). Therefore, no specific behavior can be attributed to this factor. In the MIA-exposed mice, the principal factor analysis yielded two PCs that explained 86.6% of the data set variance. PC1 represented 58.1% of the variation and included TCZ, TOZ, and %OZ (Fig. 7D and 7E). These variables relate to anxious-type behaviors in rodents. PC2 explained 38.1% of the variance and was loaded by the average and maximum speed (AS and MS, respectively, Fig. 7F), two measures associated with motor activity in mice. These data indicate that the MIA-exposed mice exhibited anxious behavior compared to the control animals.

### MIA-exposed mice show social behavior deficits

In the test for sociability, the principal component analysis yielded two factors that explain 85.1% of the data set variance for the controls (Table 1.3 and Fig. 8B). In the MIA-exposed mice, only one component was obtained, accounting for 77.0% of the variance (Table 1.3 and Fig. 8E). PC1 of the control represented 59.1% of the variance and was loaded by time investigating object (TO), number of interactions with object (NO), and total investigation time (TI) (Figs. 8B and 8C). This set of variables is related to an explorative behavior or innate curiosity of rodents. The PC2 accounted for 26.1% of the variance and was mainly loaded by socialization percentage (%S) (Figs. 8B and 8D). Although only the %S variable meets the criteria established for PC2 component loading, the contribution graph (Fig. 8D) shows that variables such as the time investigating stranger 1 (TS1) and the number of interactions with stranger 1 (NS1) also contributed to PC2. These measurements are classically associated with social interactions. In the case of MIA-exposed mice, no independent components were distinguishable, and all the measurements were loaded to PC1 (Table 1.3). These results suggest that social and exploratory behavior were different between the groups.

For the social novelty preference test, the factor analysis yielded two PCs that explained 96.7% of the variance in the data (Table 1.4 and Fig. 9B). PC1 accounted for 59.6% of the variance (Fig. 9B) and included the key related to social novelty behaviors: time investigating the stranger 2 (TS2), number of interactions with the stranger 2 (NS2), total time investigating (TI), and percent social novelty (%SN) (Fig. 9C). PC2 (37.1% variance) was mainly loaded by two variables related to social familiarity behavior: time investigating stranger 1 (TS1) and the number of interactions with stranger 1 (NS1) (Figs. 9C and 9D). For the MIA-exposed mice, two factors were obtained, PC1 representing 75.5% of the variance and PC2 21.8% (Fig. 9E). PC1 was explained by most of the measurements except for TI, which was part of PC2. Although both experimental groups presented the same number of PCs, the MIA-exposed group was made up of variables that were not related to each other, suggesting an impairment in social behaviors (Figs. 9F and 9G).

**Fig. 6.**
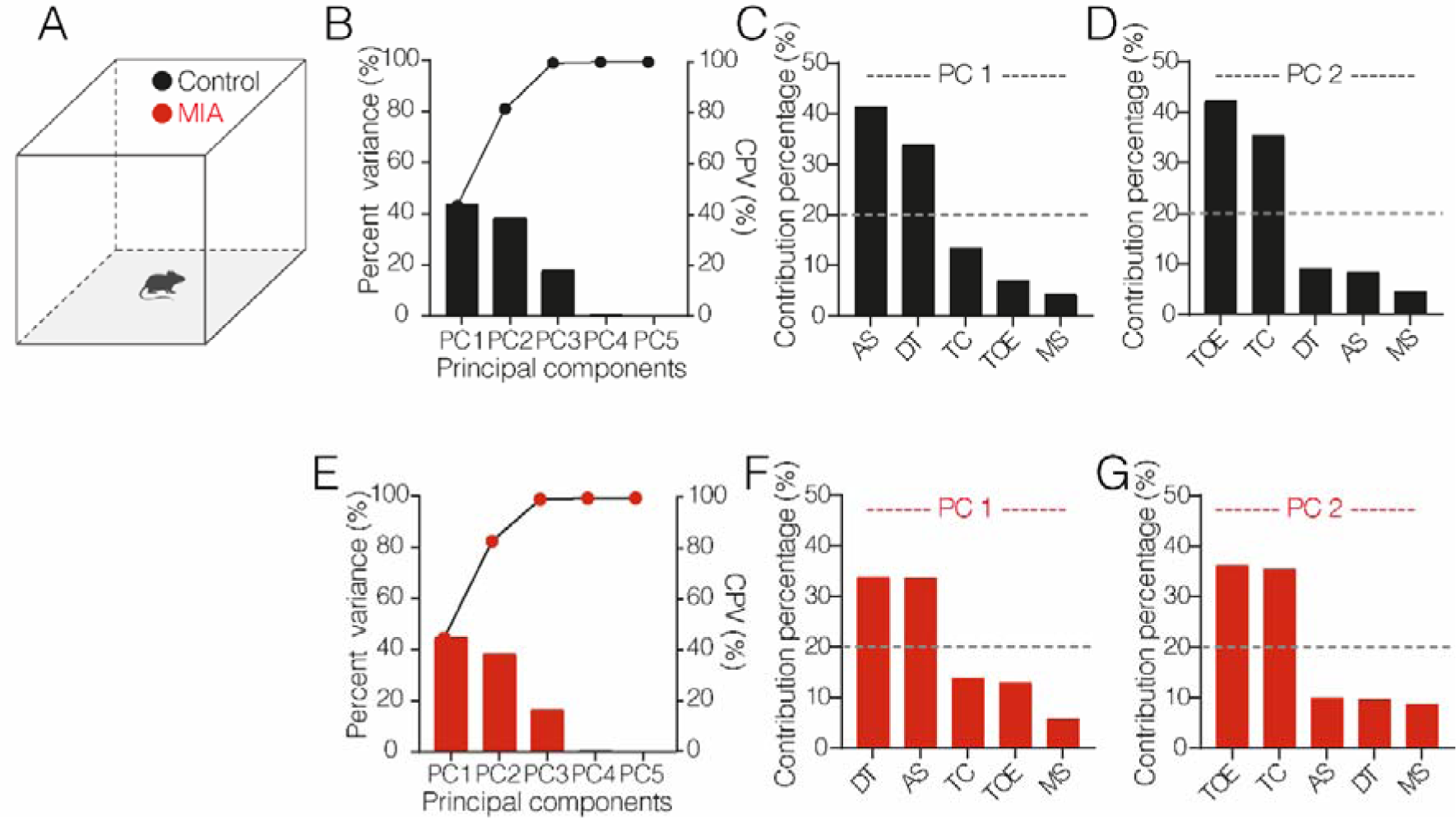
MIA alters behavioral components of the offspring in the open-field test. **A)** Schematic representation of the open-field test. **B)** Representation of the percentage of variance corresponding to each principal component (PC); the scatter plot represents the cumulative percentage of variance in the control condition. **C, D)** Bar graphs showing the percent contribution of each behavior to PC1 and PC2 in the control group. **E)** Percentage of variance corresponding to each PC and the cumulative percentage of variance in the MIA group (bars and scatter plot, respectively). **F, G)** Bar graphs showing the percent contribution of each behavior to PC1 and PC2 in the MIA group. PC1 and PC2 described >80% of the total variance in the data set in both experimental conditions. The bars that exceed the dashed lines are the behaviors that explained the principal components (C, D, F, and G). In both groups, AS and DT explained PC1, while PC2 was explained by TOE and TC. Each condition comprised eight animals from four litters.

**Fig. 7.**
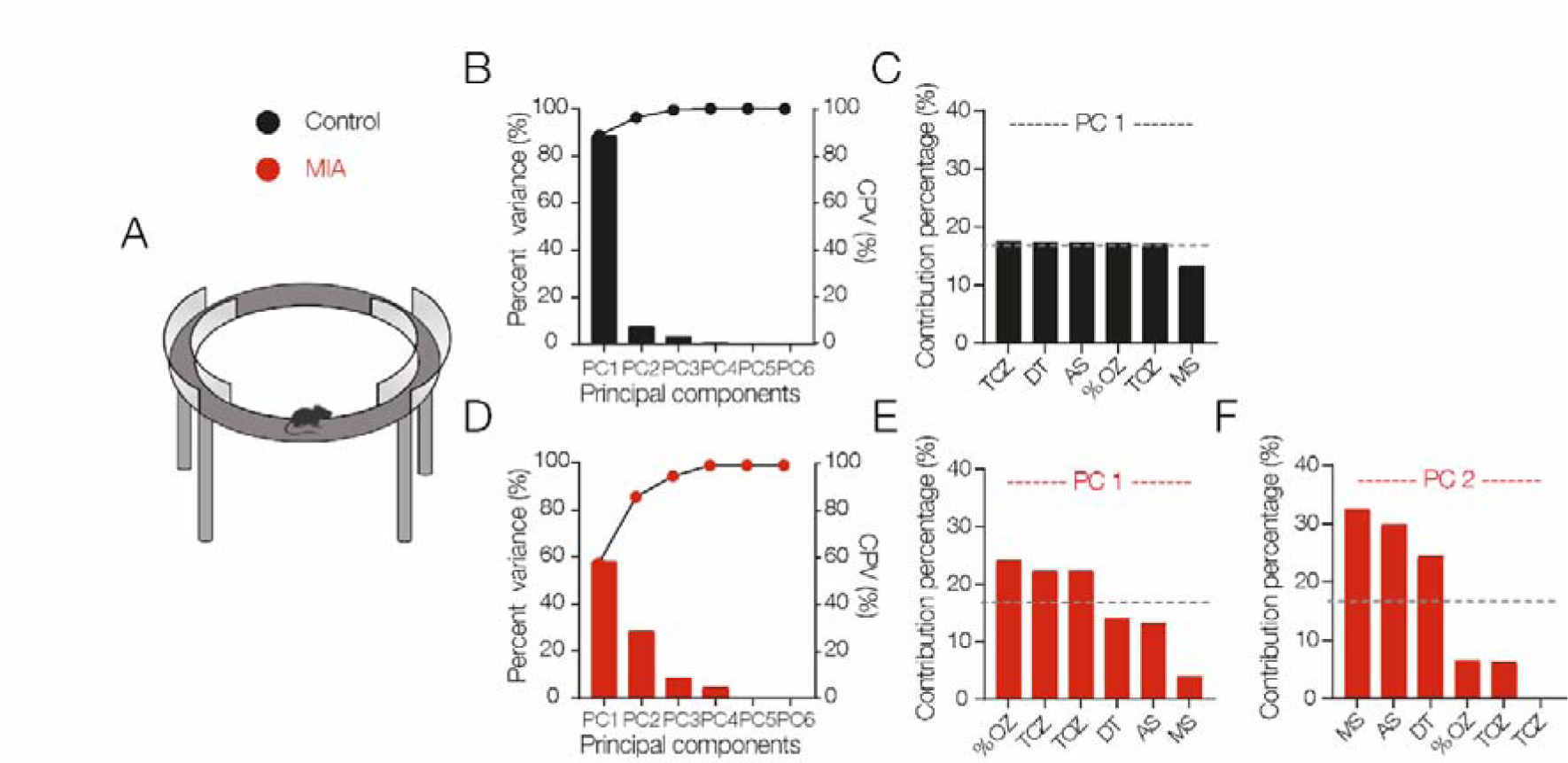
MIA alters the anxiety-like behaviors in offspring. **A)** Schematic representation of the elevated zero maze (EZM) to assess anxiety-like behavior. **B)** Bar graphs showing the percentage of variance of each PC; the scatter plot shows the cumulative percentage of variance in the control group. **C)** In the control group, PC1 was constructed by TCZ, DT, AS, %OZ, and TOZ and explained >80% of the model variance. **D)** Bar graph: the sum of PC1 and PC2 explained >80% of the total variance in the data set. **E)** Bars showing the behaviors included in PC1: %OZ, TCZ, and TOZ. For PC2, MS, AS, and DT were the main components. Dashed lines indicate the behaviors that explained the PC. Each condition comprised eight animals from four litters.

**Fig. 8.**
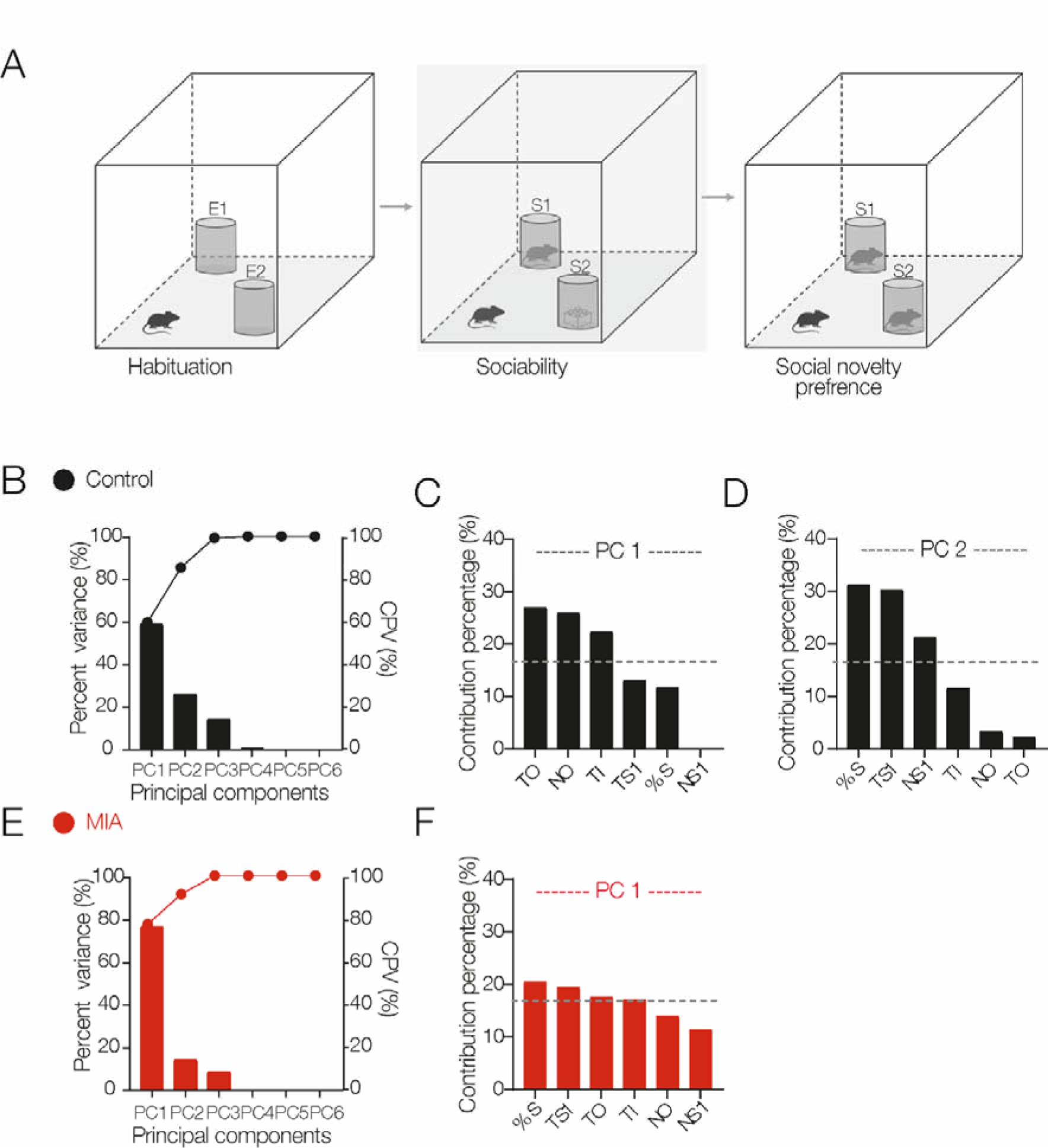
MIA impairs the sociability behavior in the offspring. **A)** Schematic representation of the modified three-chamber paradigm to assess sociability and social novelty preference, emphasizing the sociability stage. **B)** Bar graph showing the percentage of variance of each PC and the cumulative percentage of variance (scatter graph). PC1 and PC2 explained >80% of the total variance in the control animals. **C)** Bar graph. TO, NO, and TI explained PC1, whereas **D)** %S, TS1, and NS1 described PC2. **E)** Bar graph. For MIA-exposed offspring, PC1 explained >80% of the total variance. **F)** Bar graph. PC2 included %S, TS1, TO, and TI. Dashed lines represent the behaviors that explained the principal components (C, D, and F). For MIA groups, n = 5 animals / 3 litters, and for control groups, n = 5 animals / 4 litters.

**Fig. 9.**
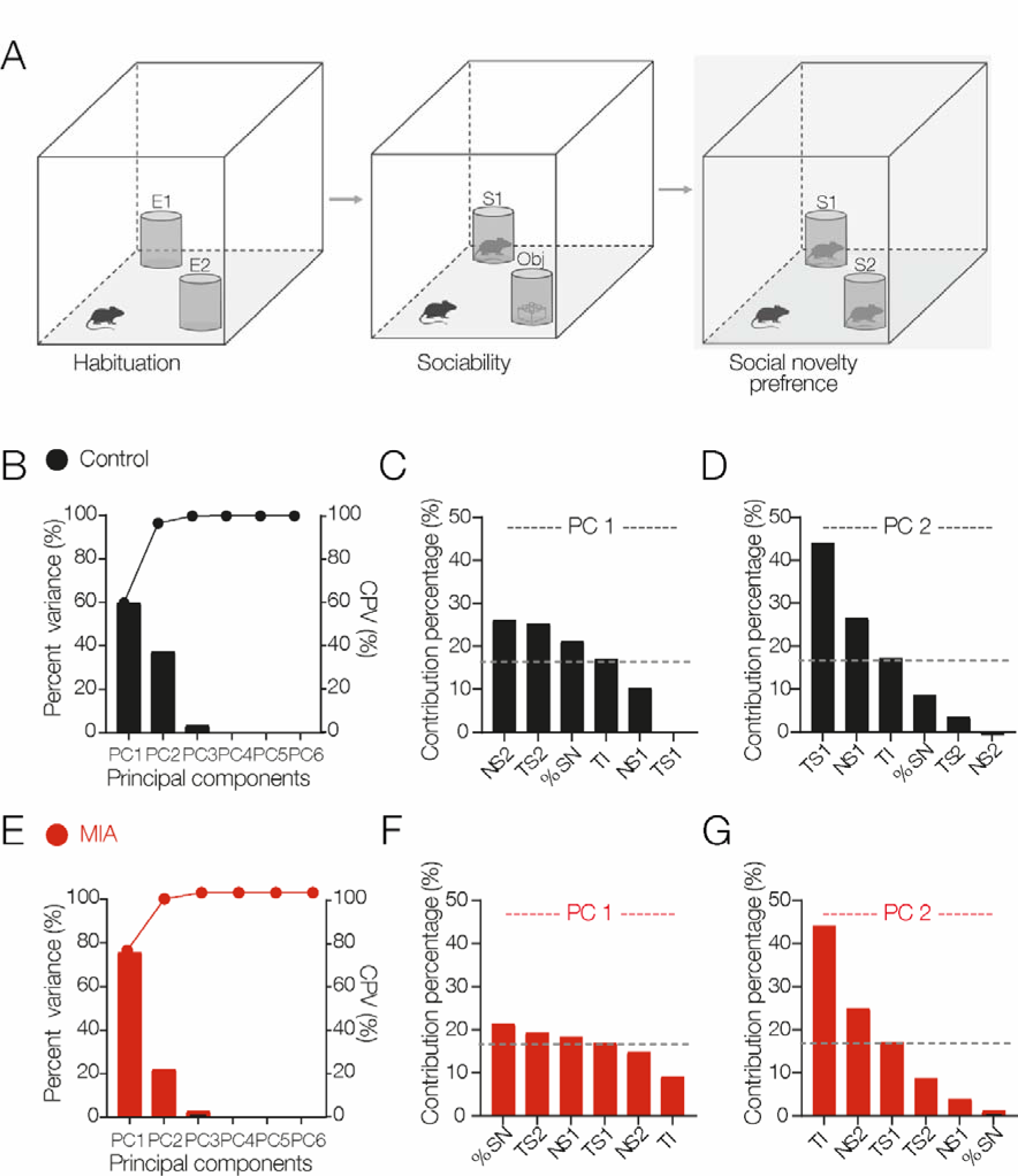
MIA alters the social novelty preference. **A)** Schematic representation of the social novelty preference stage with the modified three-chamber paradigm to assess sociability and social novelty preference. **B)** Bar graph showing the percentage of variance of each PC and the cumulative percentage of variance (scatter graph) in the control group. PC1 and PC2 explained >80% of the total variance in the control animals. **C)** Bar graph. NS2, TS2, %SN, and TI explained PC1. **D)** Bar graph. PC2 was composed of TSI, NS1, AND TI. **E)** Bar graph. For MIA-exposed offspring, PC1 and PC2 explained >80% of the total variance. **F)** Bar graph. The behaviors included in PC1 were %SN, TS2, NS1, and TS1. **G)** Bar graph. TI, NS2, and TS1 explained PC2. Dashed lines represent the behaviors that explained the principal components (C, D, F, and G). For MIA groups, n = 5 animals / 3 litters, and for control groups, n = 5 animals / 4 litters.

## 4. Discussion

The experimental evidence of this study demonstrates a series of neurophysiological alterations in GABAergic transmission that alter the computational capabilities of CA1 pyramidal cells of the dorsal hippocampus. Consistent with these findings, we have also demonstrated that animals exposed to MIA exhibit increased anxiety levels and impairments in social performance. In a previous study, we reported that the same MIA model increases intrinsic excitability and promotes a reduction in the dendritic complexity of CA1 PCs of the offspring (Griego et al., 2022). Altogether, our findings reveal molecular, cellular, and synaptic mechanisms potentially underlying the higher risk of MIA-exposed offspring developing neuropsychiatric disorders (see the proposed model, Fig. 10). To the best of our knowledge, this is the first study demonstrating that bacterial-mediated MIA alters the integration of excitatory inputs and the firing temporal fidelity in hippocampal neurons of the offspring.

**Fig. 10.**
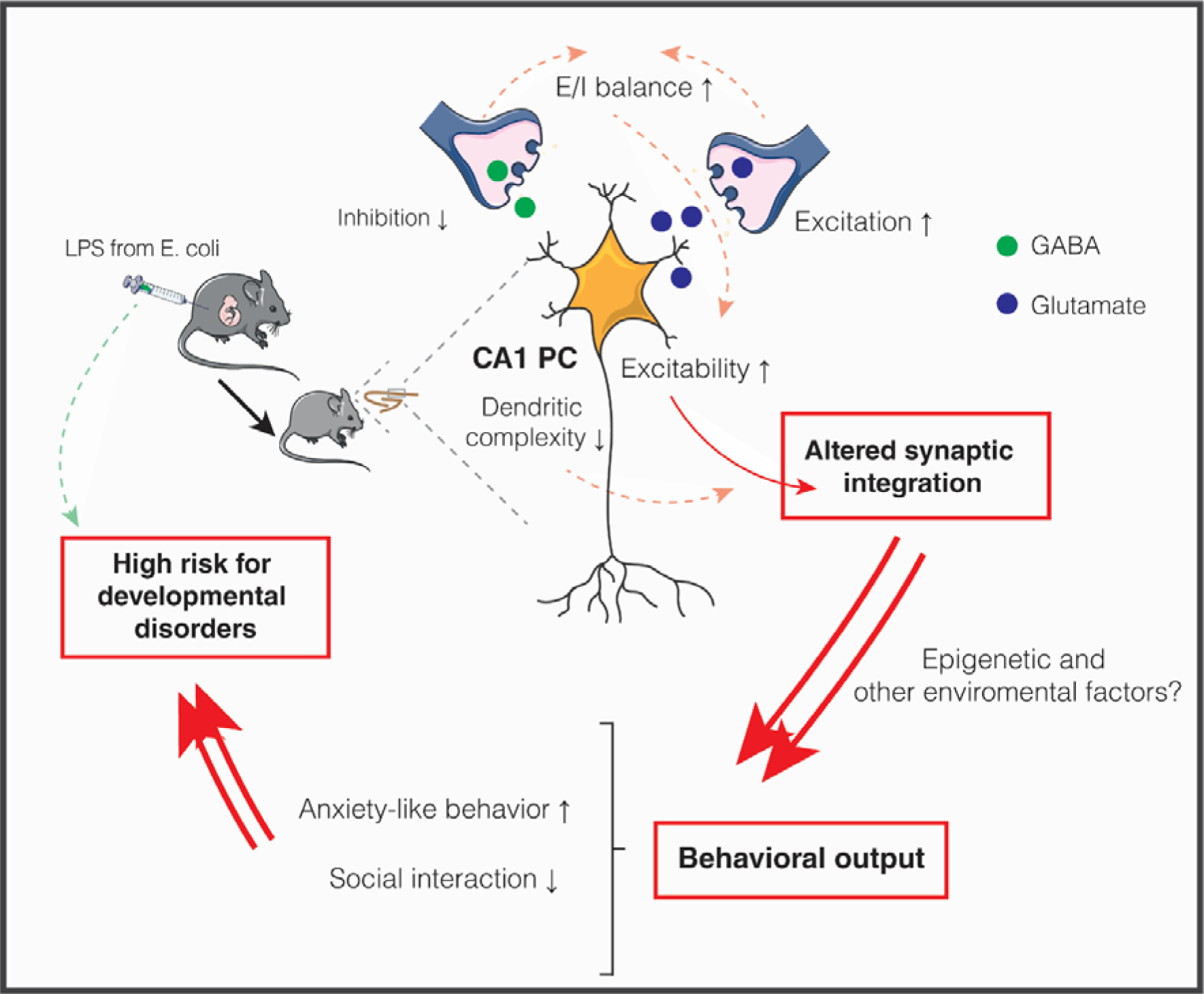
Cellular and synaptic alterations induced by MIA in CA1 PC of the offspring. Illustrative cartoon showing that maternal inflammation induced by LPS injection leads to neurodevelopmental impairments in the offspring, which in turn result in several alterations in CA1 PC physiology, such as a reduction in morphological complexity, a shift toward excitation of the E-I balance, and increased somatic excitability. Altogether, these cellular and synaptic changes impair the computational capabilities of the CA1 PC, leading to abnormal synaptic integration and a loss of spike generation temporal fidelity. All these neurophysiological alterations, in combination with epigenetic and other environmental factors, may underlie the behavioral phenotype observed in this model, which is characterized by impaired performance in social interaction as well as an increase in anxiety-like behavior.

### 4.1 Changes in synaptic integration and GABAergic inhibition

Dendritic integration represents the phenomenon by which neurons transform excitatory inputs into a scaled output, as a general rule, in the form of an action potential (Gulledge et al., 2005). This computational phenomenon depends primarily on three factors: biophysical membrane properties, dendritic geometric properties, and GABAergic inhibition, especially at the perisomatic level (Freund & Katona, 2007; Magee, 2000; Spruston, 2008). In that sense, we found that both temporal and spatial components of synaptic integration were facilitated in the CA1 PCs of MIA-exposed mice. On the one hand, temporal summation enables the computation of successive excitatory inputs from the same presynaptic origin (Hao et al., 2009) that may result in an action potential or the attenuation of the repetitive excitatory drive. On the other hand, spatial summation enables the integration of excitatory inputs coming from different sources (Hao et al., 2009). We previously reported that LPS-induced MIA increases the R_N_ and τ_memb_ of the CA1 PCs of the offspring (Griego et al., 2022), membrane properties on which dendritic integration also relies. More importantly, both components of dendritic integration are highly sensitive to changes in GABAergic inhibition. Consistent with this idea, a previous study has shown that perisomatic inhibition frames the temporal fidelity in such a way that two excitatory inputs converging within this frame are enhanced with such efficiency that they generate an action potential (Pouille & Scanziani, 2001).

Our results show that MIA causes a decrease in the number of GAD67-expressing cells and induces a disequilibrium in the E-I balance, moving it toward a hyperexcitable state. Notably, we also noticed that extracellular GABA receptors appear not to have played a major role, as GABAergic I_tonic_ was unchanged in MIA-exposed mice. In this regard, a consequence commonly reported in different models of MIA is the appearance of qualitative and quantitative alterations in GABAergic interneurons, both in the cerebral cortex and the hippocampus of the offspring, especially PV-IN (Canetta et al., 2016; Nakamura et al., 2021; Nakamura et al., 2019; Vojtechova et al., 2021). Surprisingly, we found a decrease in WIN-sensitive inhibition but not in DAMGO-sensitive inhibition, which may be indicative of alterations in GABAergic interneurons expressing CB1R but not MOR (CCK-IN and PV-IN, respectively). These findings are particularly important given that CCK-IN and PV-IN provide perisomatic inhibition on CA1 PCs (Freund & Katona, 2007), with this decrease in inhibition coming from CCK-IN being a potential and very likely mechanism for the facilitation in the temporal and spatial summation induced by MIA. GABAergic transmission, local inhibitory circuits, and all the molecular machinery associated with maintaining the E-I balance undergo a substantial maturation process in prenatal and early postnatal stages (Di Cristo, 2007; Lopatina et al., 2019). Importantly, the molecular mechanisms regulating this maturation are sensitive to the action of molecules of immunity, such as the activation of Toll-like receptors (TLRs) and several cytokines whose release is a well-documented consequence of MIA (Canales et al., 2021; Fernandez et al., 2019; Griego et al., 2022). Data reported in this study strongly suggest that LPS-induced MIA promotes developmental impairments in the mechanisms governing the integration of interneurons into neural circuits and, in turn, an imbalance in the E-I ratio that eventually leads to an aberrant dendritic integration of synaptic inputs onto CA1 PCs. All this is in addition to the impairments in the intrinsic excitability and dendritic morphology previously reported (Griego et al., 2022).

### 4.2 Anxiety-like behavior and deficits in the social performance of the MIA-exposed offspring

The hippocampal circuits have been proven to be neural substrates for learning and memory (Basu & Siegelbaum, 2015; Jarrard, 1993; Whitlock et al., 2006). In consequence, perturbations in the anatomical and functional properties of the hippocampus are the signature of several neuropsychiatric and neurologic disorders affecting cognitive execution (Banker et al., 2021; Busche et al., 2012; Harrison, 2004). In this work, we complemented cellular and synaptic investigations with behavioral analysis of the MIA-exposed offspring. Our data show a significant impairment in social behavior and increased anxiety-like behaviors in the offspring. These behavioral findings are consistent with the behavioral alterations associated with the neurodevelopmental disorders to which MIA has been linked, including schizophrenia and ASD (Dodell-Feder et al., 2015; Pinkham et al., 2020). Previous studies have also documented that MIA promotes deficiencies in spatial memory (Murray et al., 2017; Wolff et al., 2011), as well as in sensorimotor and social performance (Choi et al., 2016; Glass et al., 2019; Gzieło et al., 2023; Mueller et al., 2021). It is feasible to hypothesize that MIA-induced alterations in offsprings’ hippocampus structure and function are underlying the reported behavioral disturbances. However, causal studies linking neurophysiological alterations to the behavioral phenotype described here and by others require further investigation.

### 4.3 Limitations of this study and future directions

Since sex and gender bias in experimental neuroscience represents a tangible problem in biomedical research (Coiro & Pollak, 2019), a caveat of our study is that the experiments were restricted to male mice. Nonetheless, electrophysiological recording in post-pubertal female mice is challenging due to fluctuations in physiologic and behavioral responses depending on the specific phase of the estrous cycle (Chari et al., 2020; Lovick & Zangrossi, 2021; Rocks & Kundakovic, 2023; Scharfman et al., 2003). In that regard, a consideration of that magnitude is beyond this manuscript’s objective. However, some previous studies have reported that male and female offspring from dams injected with LPS or Poly I:C during mid-gestation show dissimilar behavioral phenotypes, suggesting that females are better protected against developmental damages leading to neuropsychiatric-like alterations in rodents (Haida et al., 2019; Xuan & Hampson, 2014). This is of paramount relevance since ASD, one of the neurodevelopmental disorders most commonly associated with MIA, affects three males for every one female (Loomes et al., 2017). The mechanistic explanation for this male bias is not well known, but potential mechanisms may include hormonal and genetic influences.

Notwithstanding the difficulty in establishing such a causal connection, efforts focused on updating and improving experimental designs, starting by considering sex and gender as indisputable biological variables, are of prime importance.

## 5. Conclusions

The exposition to bacterial antigens induces an intense maternal immune response that impairs fetal neurodevelopment. The data presented in this study provide an important update of the neurophysiological alterations related to inflammatory insults during mid-gestation. This study shows, for the first time, that the modeling of bacterial infections in pregnant mice leads to significant changes in synaptic summation and spike generation temporal fidelity of hippocampal neurons, which in turn might act as a substrate for behavioral disturbances in the juvenile offspring. These results provide novel mechanisms whereby maternal immune system activity may contribute to the development of ASD and other MIA-related neuropsychiatric disorders in the offspring.

## Acknowledgments

The authors wish to thank Juan Javier López for technical assistance. Acknowledgments are extensive to Dr. Claudia González-Espinosa and Dr. Deisy Segura-Villalobos for generous support in performing ELISA assays. Some of the experiments included in this manuscript are part of the doctoral thesis of Dr. Ernesto Griego.

---

**Supplementary Fig. S1**. Effects of MIA on the temporal summation of Schaffer collateral inputs at 20 Hz. **A)** Representative traces of the EPSP trains at 20 Hz in both experimental conditions. **B)** Scatter plot showing EPSP amplitudes along the trains evoked by Schaffer collateral stimulation at 20 Hz. **D)** Violin plot summarizing the lack of effect of MIA on the temporal summation of EPSPs evoked at 20 Hz in the CA1 PCs of the offspring.

**Table 1.1.**
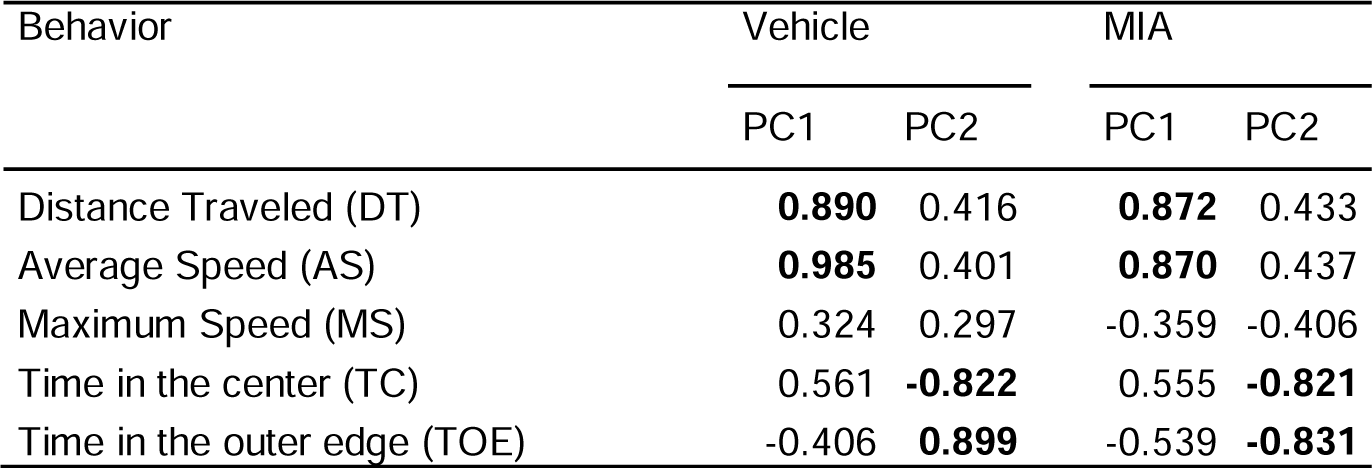
Factor loadings of open field test behaviors of both VEH and LPS-treated animals.

**Table 1.2.** Factor loadings of elevated zero maze behaviors of both VEH and LPS-treated animals.

| Behavior | Vehicle |  | MIA |  |
| --- | --- | --- | --- | --- |
|  | PC1 |  | PC1 | PC2 |
| Distance Traveled (DT) | **0.961** |  | 0.700 | 0.641 |
| Average Speed (AS) | **0.960** |  | 0.679 | **0.709** |
| Maximum Speed (MS) | **0.841** |  | 0.372 | **0.739** |
| Time in the closed zone (TCZ) | **-0.966** |  | **-0.881** | 0.046 |
| Time in the opened zone (TOZ) | **0.952** |  | **0.881** | -0.327 |
| Percent time in the opened zone (% OZ) | **0.957** |  | **0.918** | -0.331 |

**Table 1.3.** Factor loadings of sociability test behaviors of both VEH and LPS-treated animals.

| Behavior | Vehicle |  | MIA |  |
| --- | --- | --- | --- | --- |
|  | PC1 | PC2 | PC1 | PC2 |
| Time investigating Strange 1 (TS1) | 0.680 | 0.688 | **0.946** |  |
| No. of interactions Stranger 1 (NS1) | 0.004 | 0.577 | **0.724** |  |
| Time investigating Object (TO) | **0.979** | -0.185 | **-0.903** |  |
| No. of interactions Object (NO) | **0.958** | -0.229 | **-0.804** |  |
| Total Investigation Time (TI) | **0.89** | 0.427 | **0.889** |  |
| Socialization percentage (% S) | -0.644 | **0.7** | **0.9ta74** |  |

**Table 1.4.** Factor loadings of social novelty preference test behaviors of both VEH and LPS-treated animals.

| Behavior | Vehicle |  | MIA |  |
| --- | --- | --- | --- | --- |
|  | PC1 | PC2 | PC1 | PC2 |
| Time investigating Strange 1 (TS1) | 0.064 | **0.989** | **-0.877** | 0.474 |
| No. of interactions Stranger 1 (NS1) | -0.605 | **0.768** | **-0.911** | -0.227 |
| Time investigating Strange 2 (TS2) | **0.95** | 0.277 | **0.935** | 0.338 |
| No. of interactions Strange 2 (NS2) | **0.967** | 0.037 | **0.82** | -0.571 |
| Total Investigation Time (TI) | **0.782** | 0.62 | 0.643 | **0.76** |
| Social Novelty percentage (%) (% SN) | **0.871** | -0.439 | **0.986** | -0.129 |

